# The opposite aging effect to single cell transcriptome profile among cell subsets

**DOI:** 10.1101/2024.05.01.591990

**Authors:** Daigo Okada

**Affiliations:** Center for Genomic Medicine, Graduate School of Medicine, Kyoto University, Kyoto 606-8507, Japan

## Abstract

Comparing transcriptome profiling between younger and older samples reveals genes related to aging and provides insight into the biological functions affected by aging. Recent research has identified sex, tissue, and cell type-specific age-related changes in gene expression. This study reports the overall picture of the opposite aging effect, in which aging increases gene expression in one cell subset and decreases it in another cell subset. Using the Tabula Muris Senes dataset, a large public single-cell RNA sequencing dataset from mice, we compared the effects of aging in different cell subsets. As a result, the opposite aging effect was observed widely in genome-wide genes, particularly enriched in genes related to ribosomal function and translation. The opposite aging effect was observed in the known aging-related genes. Furthermore, the opposite aging effect was observed in the transcriptome diversity quantified by the number of expressed genes and the Shannon entropy. This study highlights the importance of considering the cell subset when intervening with aging-related genes.

## Introduction

Comparing transcriptome profiles between younger and older samples is a powerful approach to identifying genes associated with aging. Comprehensive identification of age-related changes in gene expression can provide a detailed description of biological functions that are altered by aging. While changes in gene expression that are common to all cellular aging have been studied [9, 22, 30], sex [1, 12], tissue [25], and cell type [7, 30] specific changes in aging-related gene expression have been identified.

Recent studies on expression quantitative trait loci (eQTL) have reported cases in which the effects of genetic factors on gene expression act in opposite directions in different cell subsets. For example, the eQTL with opposite positive and negative effects on gene expression levels between different tissues has been reported and called the opposite eQTL effect [16]. The integrated analyses of single-cell RNA-seq (scRNA-seq) and genomic data have also identified eQTLs with opposite effects on different cell types [19, 29]. These findings suggest that the effect of genetic perturbation on the transcriptome is cell subset-dependent.

In this study, we analyzed a large public dataset of mouse scRNA-seq and reported the landscape of opposite effects of aging among cell subsets on the transcriptome. The report of this study highlights the importance of considering the cell subsets when intervening with aging-related genes.

## Method

### Data and preprocessing

We analyzed data from the Tabula Muris Senes dataset, which is a large public single-cell RNA-seq dataset from mice of different ages [5]. We used the TabulaMurisSenisData package in R to obtain pre-processed data after log transformation [23]. The Tabula Muris Senes project provided two different mouse single-cell RNA-seq datasets from different experimental designs (FACS or droplet). FACS dataset contains the gene expression data of 22966 genes × 110824 cells and Droplet dataset contains that of 20138 genes × 245389 cells. We analyzed each of the two data sets to validate the stability of our results. Cells derived from 3-month-old mice were defined as the young group, and cells derived from 18-month-old or older mice were defined as the old group. In this analysis, cells derived from different cell types, sexes, and tissues were treated as different cell subsets. Cell subsets containing more than 25 cells in both young and old groups were used for subsequent analysis. Finally, 189 cell subsets in the FACS dataset and 99 cell subsets in the droplet dataset were used in this analysis. A list of cell types used can be found in Supplementary Table 1.

### Quantifying the opposite aging effect on gene expression

In each cell subset, we applied Welch’s t-test to the gene expression levels in the pre-processed scRNA-seq dataset to compare the differential expression of each gene between the old and young groups. We applied Bonferroni correction of P values to assess statistical significance and counted the number of cell subsets for which a significant increase or decrease was observed for each gene. In this step, Bonferroni correction was performed for each of the datasets, and the total number of tests is the number of all cell subsets and gene combinations.

We defined the Opposite Effect Score to detect the situation where aging increases gene expression in one cell subset and decreases it in another cell subset. Let *N*_*up*_ and *N*_*down*_ be the number of cell subsets where the expression level of a gene increased or decreased with aging, respectively. Here, the following Opposite Effect Score can be an indicator of the strength of the opposite aging effect for the gene.

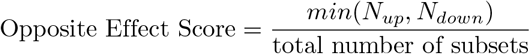

Genes for which *N*_*up*_ and *N*_*down*_ were both *>* 0 were considered as genes with the opposite aging effect. Genes for which *N*_*up*_ *>* 0 and *N*_*down*_ = 0 were considered as “UP-only genes”. Genes with *N*_*down*_ *>* 0 and *N*_*up*_ = 0 were considered “DOWN-only genes”. We performed Gene Ontology (Biological Process) enrichment analysis on genes with Opposite Effect Score *>* 0.05 using the WebGestaltR package in R [11]. The database was set to geneontology Biological Process noRedundant. The false discovery rate (FDR) threshold was set to 0.05.

### Evaluation of the opposite aging effect on transcriptome diversity

We investigated the opposite aging effect on transcriptome diversity. The simplest measure of transcriptome diversity is the number of genes expressed in a cell. How the number of expressed genes changes with age is controversial. For example, while the number of expressed genes decreases with age in mouse oocytes [3], it increases in mouse blood [2].

Another method to quantify the transcriptome diversity is Shannon entropy [14]. This method can take into account not only zero or non-zero expression of genes but also continuous expression level.

Transcriptome diversity based on Shannon entropy is calculated as follows. The normalization of the sum of expression levels to 1 can be treated as the relative frequency of transcription of genes. Let *p*_*i*_ be the relative frequency of transcription of the i-th gene in a single cell (*i* = 1, 2, ., *N*), the diversity (*H*) of the transcriptome in each tissue can be quantified by applying the Shannon entropy equation as

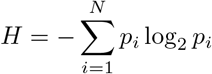

Transcriptome diversity based on Shannon entropy is associated with biological variables such as cancer [15]. It can also be used to search for optimal drug concentrations in cell culture [20]. Recently, it is also used in single-cell RNA-seq data analysis [6, 10]. Although it can be used as a good biological feature of the overall transcriptome profile, the relationship between Shannon entropy-based transcriptome diversity and aging is unclear.

We calculated the log2 of the number of expressed genes and Shannon entropy for single cells and examined their age-related change for each cell subset. We evaluated the difference in transcriptome diversity scores between the young and old groups for each cell subset using Welch’s t-test. Under the Bonferroni criterion, the threshold for the P-value was set at 0.05 / the number of subsets. We also calculated Cohen’s D, a representative effect size measure of the difference in means between groups, using the effsize package in R [27].

## Result

We analyzed the Tabula Muris Senes dataset to investigate the landscape of the opposite aging effect on gene expression among cell subsets. Figure 1A shows a scatterplot of the number of cell subsets with increased (Up) and decreased (Down) expression. While there are genes that show only up or down, we also find genes that show reversed aging changes in multiple subsets. We observed the opposite aging effect on gene expression in 15.1 % and 9.7 % of the genes in two datasets, respectively (Figure 1B). The results suggest opposite aging effect was widely observed in genome-wide gene expression profiles.

**Figure 1.**
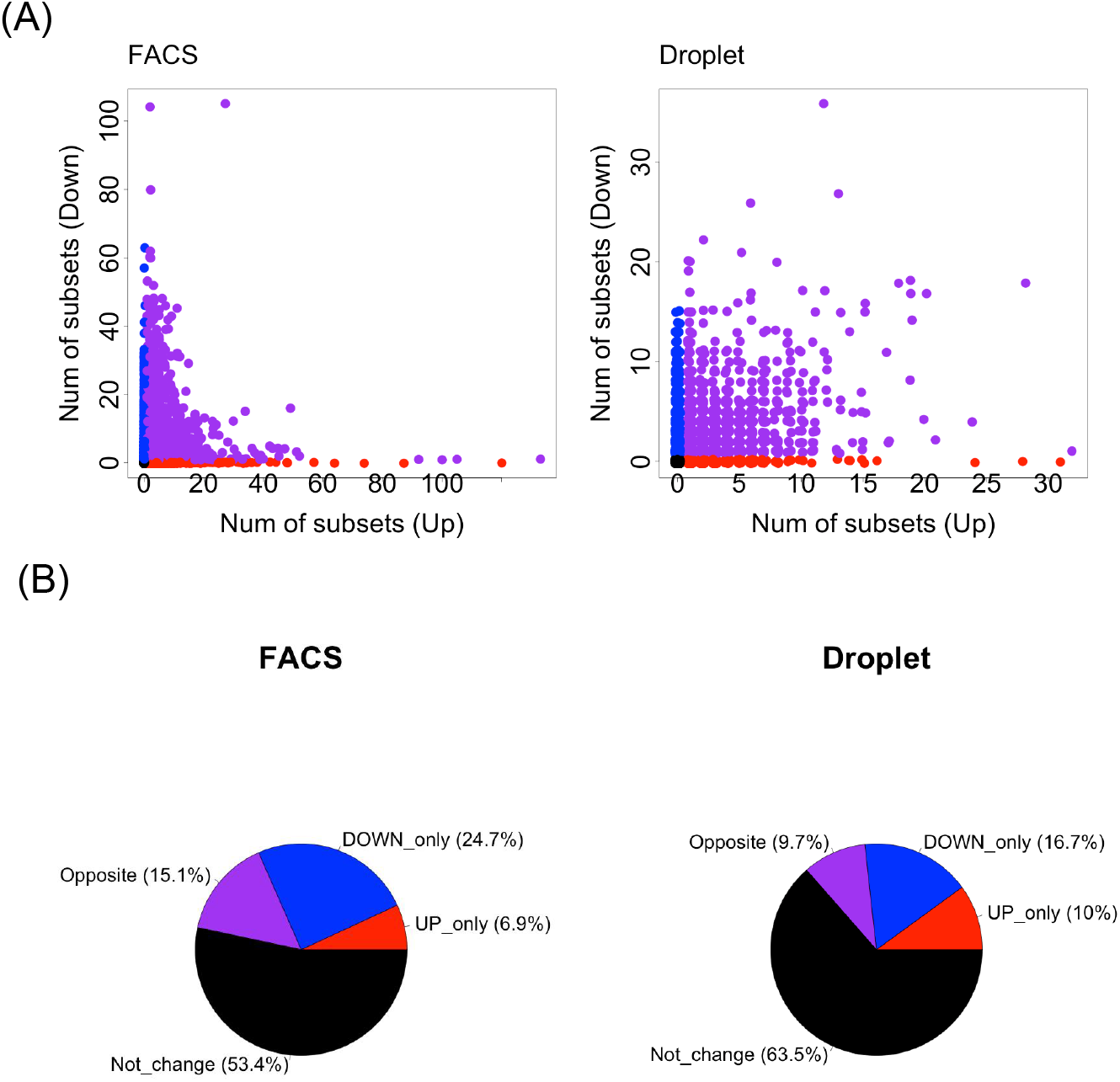
(A)Scatterplots of the number of subsets whose gene expression increased (*N*_*up*_) or decreased (*N*_*down*_) with aging for genome-wide genes. The genes of “UP only “ (*N*_*up*_ *>* 0, *N*_*down*_ = 0), “DOWN only” (*N*_*up*_ = 0, *N*_*down*_ *>* 0), “Opposite” (*N*_*up*_ *>* 0, *N*_*down*_ *>* 0), “Not change” (*N*_*up*_ = 0, *N*_*down*_ = 0) are colored by red, blue and purple, respectively. Data points were jitterd to prevent points from being plotted on top of each other. (B)Percentage of the genes with “UP only”, “DOWN only”, “Opposite”, and “Not change” among genome-wide genes.

The histogram of opposite effect scores of genome-wide genes suggests that a strong opposite aging effect is observed in some genes (Figure 2A). The genes with the top ten opposite effect scores are shown in Figure 2B. In Gene Ontology analysis, the 14 and 18 Gene ontology terms were significantly enriched (FDR *<* 0.05) (Figure 3). Among these terms, GO:0022613 (ribonucleoprotein complex biogenesis) and GO:0002181 (cytoplasmic translation) were the gene ontology terms with the top two enrichment ratio in both datasets. These results suggest that the direction of age-related change of ribosome-associated gene expression depends on cell subsets. The *N*_*up*_, *N*_*down*_, and opposite effect scores for all genes in each dataset are available in Supplementary Tables S2 and S3, respectively.

**Figure 2.**
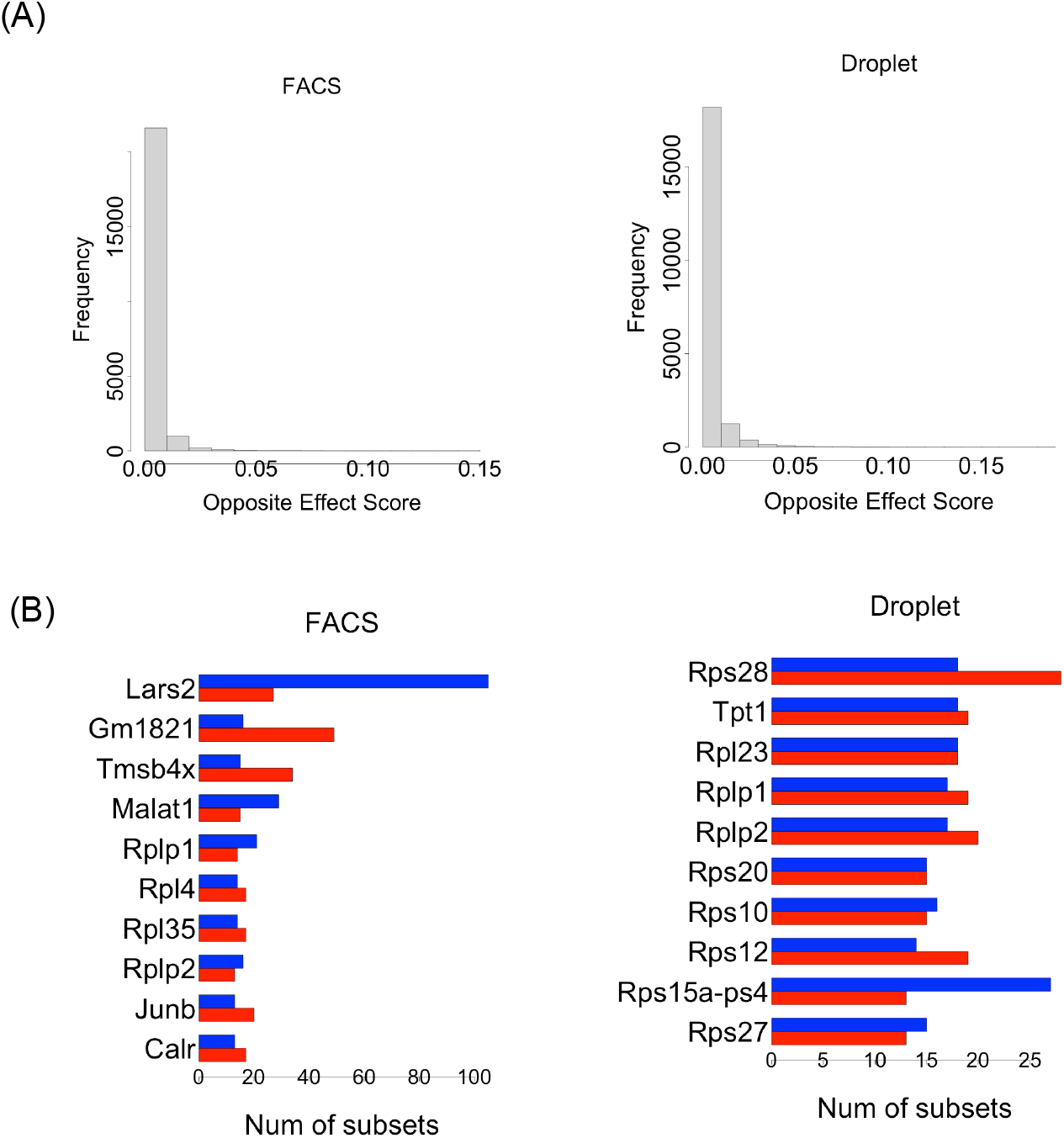
(A) Histogram of opposite effect score for genome-wide genes. (B)The genes with the top ten opposite effect scores. The red bar represents the number of subsets whose gene expression increased (*N*_*up*_). The blue bar represents the number of subsets whose gene expression decreased (*N*_*down*_).

**Figure 3.**
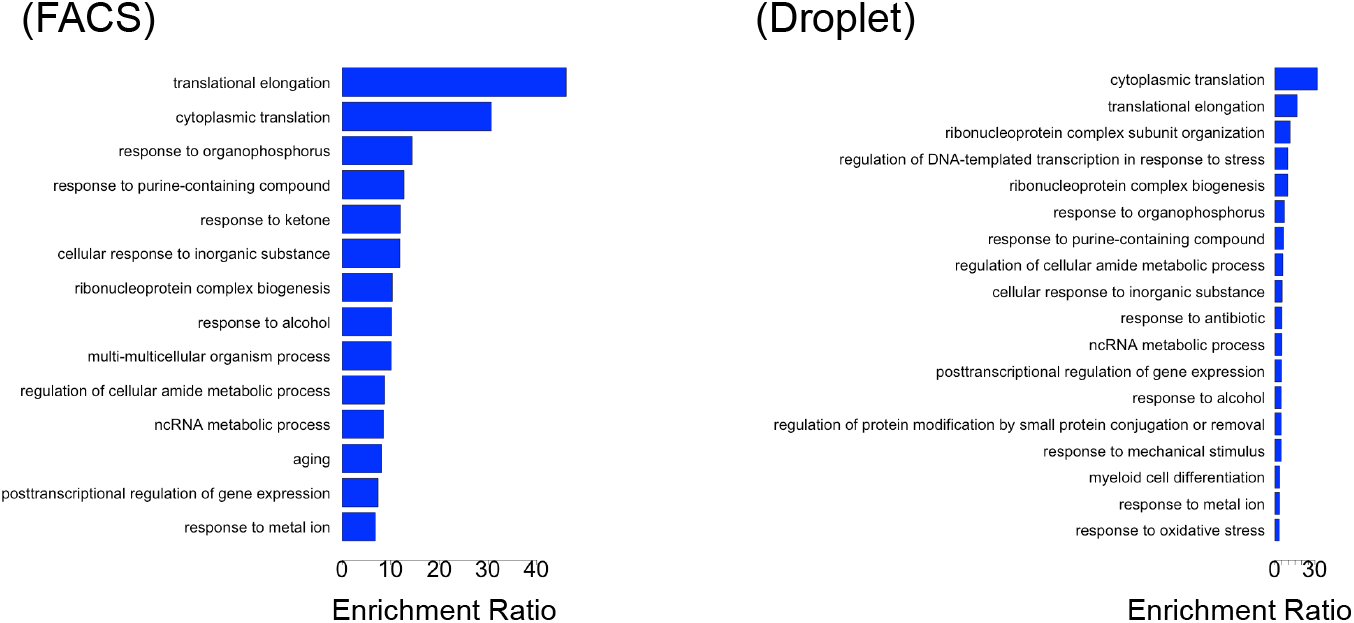
The Results of enrichment analysis to Gene Ontology (Biological Process) for the gene sets with the opposite effect score *>* 0.05. The X-axis represents the enrichment ratio of each gene ontology term.

We also examined the presence or absence of the opposite aging effect in the known 183 aging-associated genes obtained from the GenAge database [26]. In 37.5 and 21.2 % of the GenAge genes, the opposite aging effect was observed in two datasets, respectively (Figure 4A). In the 59 genes (32.2 %), the opposite aging effect was observed in at least one data set. The genes with the top ten opposite aging scores are shown in Figure 4B. *Lmna, Mt1*, and *Cebpb* were in the top ten in both data sets and their expression has been shown in Figure 5 as an example. *Lmna* encodes a nuclear envelope protein involved in maintaining nuclear structure and regulating transcription and is known to be related to aging and longevity [8, 13]. *Mt1* is a gene involved in metal ion metabolism, and its cardiac-specific over-expression extends lifespan [28].*Cebpb* is a gene involved in fat metabolism, and replacement of the *Cebpa* gene with *Cebpb* is reported to extend life span [4].

**Figure 4.**
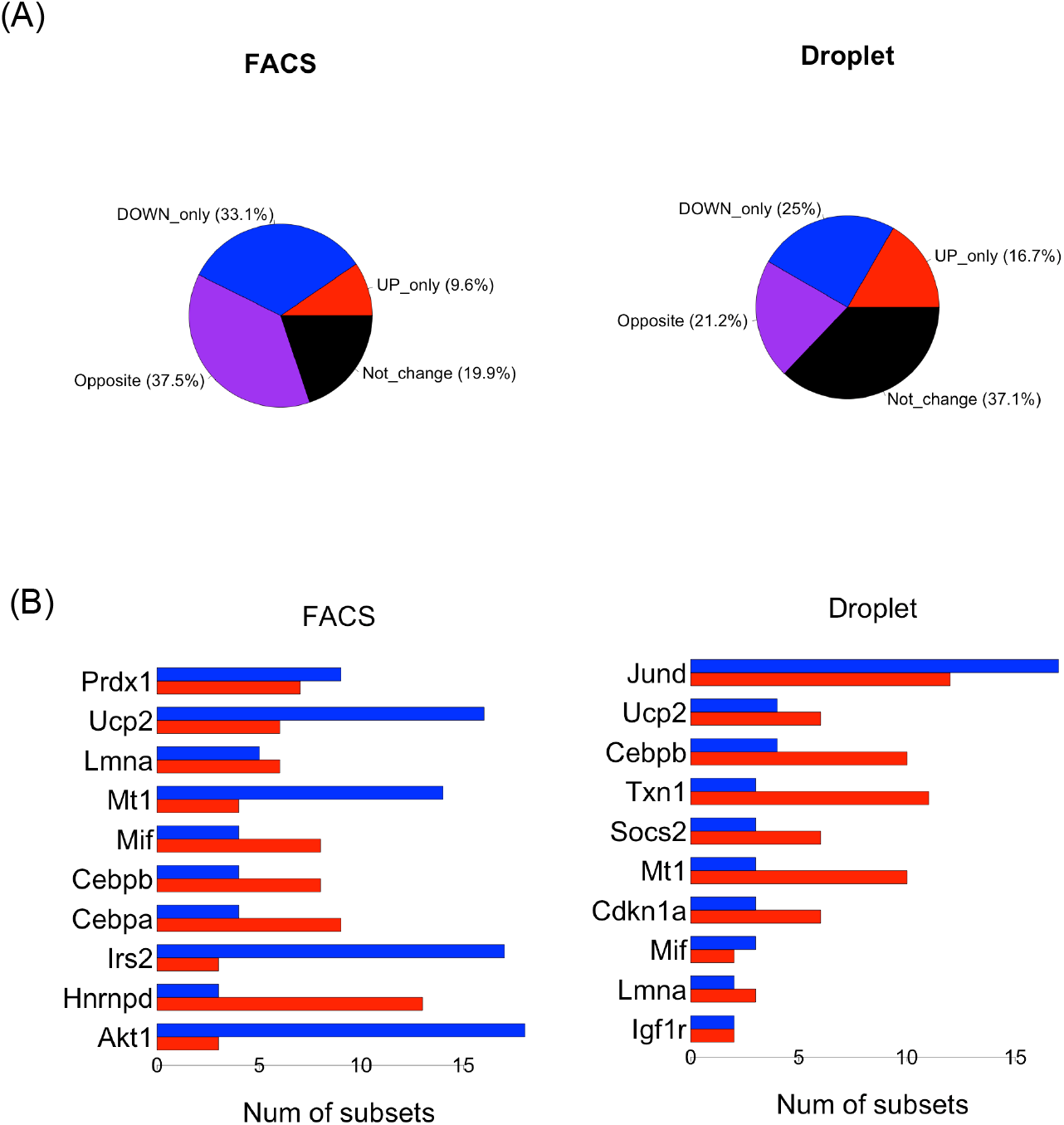
opposite aging effect on the known mouse age-related genes registered in GenAge database. (A)Percentage of the genes with “Up only” (*N*_*up*_ *>* 0, *N*_*down*_ = 0), “Down only” (*N*_*up*_ = 0, *N*_*down*_ *>* 0), “Opposite” (*N*_*up*_ *>* 0, *N*_*down*_ *>* 0), and “Not change” (*N*_*up*_ = 0, *N*_*down*_ = 0) among GenAge genes. (B) The GenAge genes with the top ten opposite effect scores.The red bar represents the number of subsets whose gene expression increased (*N*_*up*_). The blue bar represents the number of subsets whose gene expression decreased (*N*_*down*_).

**Figure 5.**
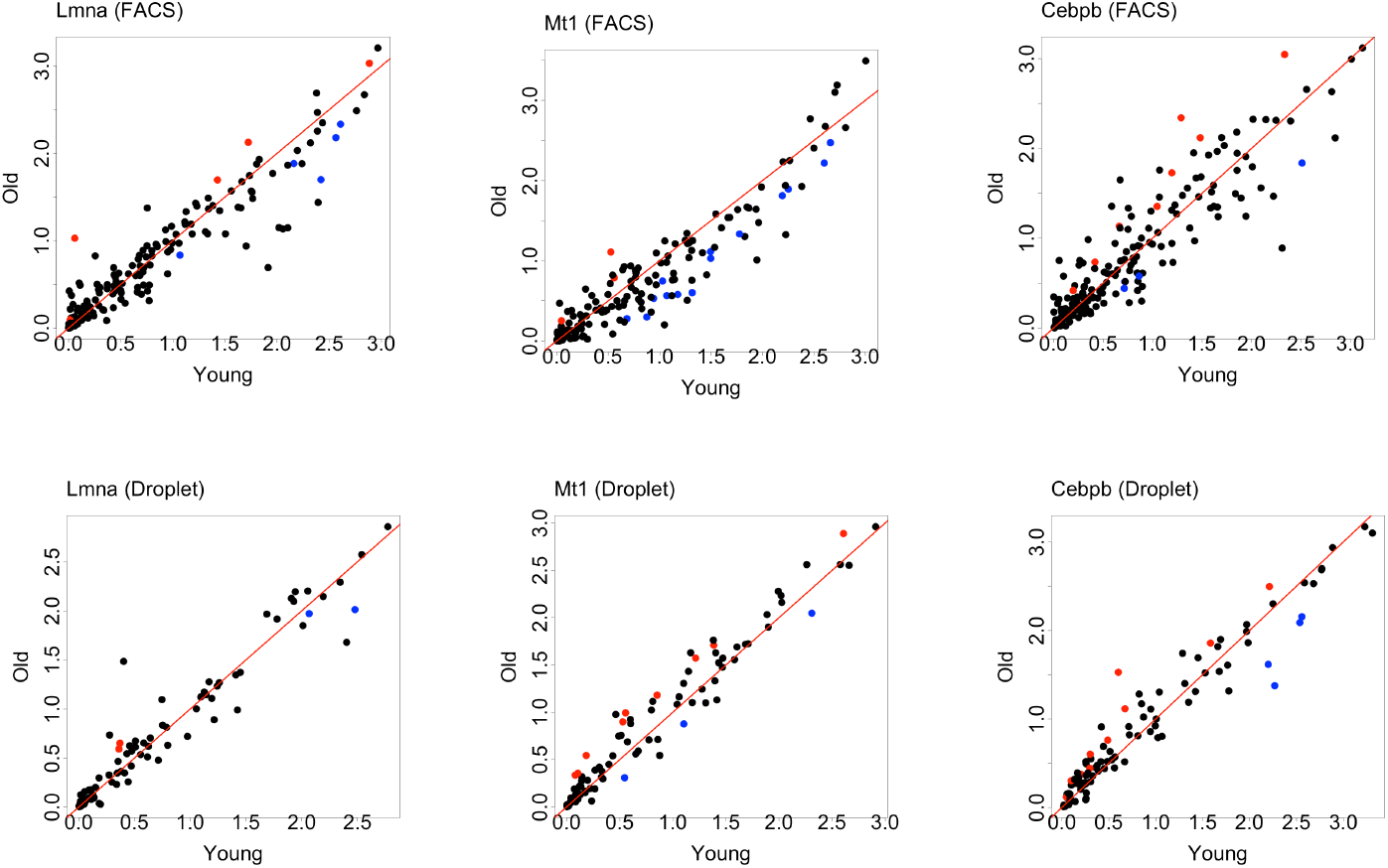
Scatterplots of mean expression in the young group (x-axis) and old group (y-axis) for *Lmna, Mt1*, and *Cebpb* in the all cell subsets. Cell subsets with statistically significant differences identified are plotted in red (Up) or blue (Down). The red line represents y=x.

Figure 6 shows a volcano plot of age-related change in transcriptome diversity. The opposite aging effect was observed in transcriptome diversity. The number of cell subsets with decreasing transcriptome diversity with aging was greater than that of the subset with increasing transcriptome diversity. This observation is true for both FACS and droplet datasets and on both the number of expressed genes and Shannon entropy.

**Figure 6.**
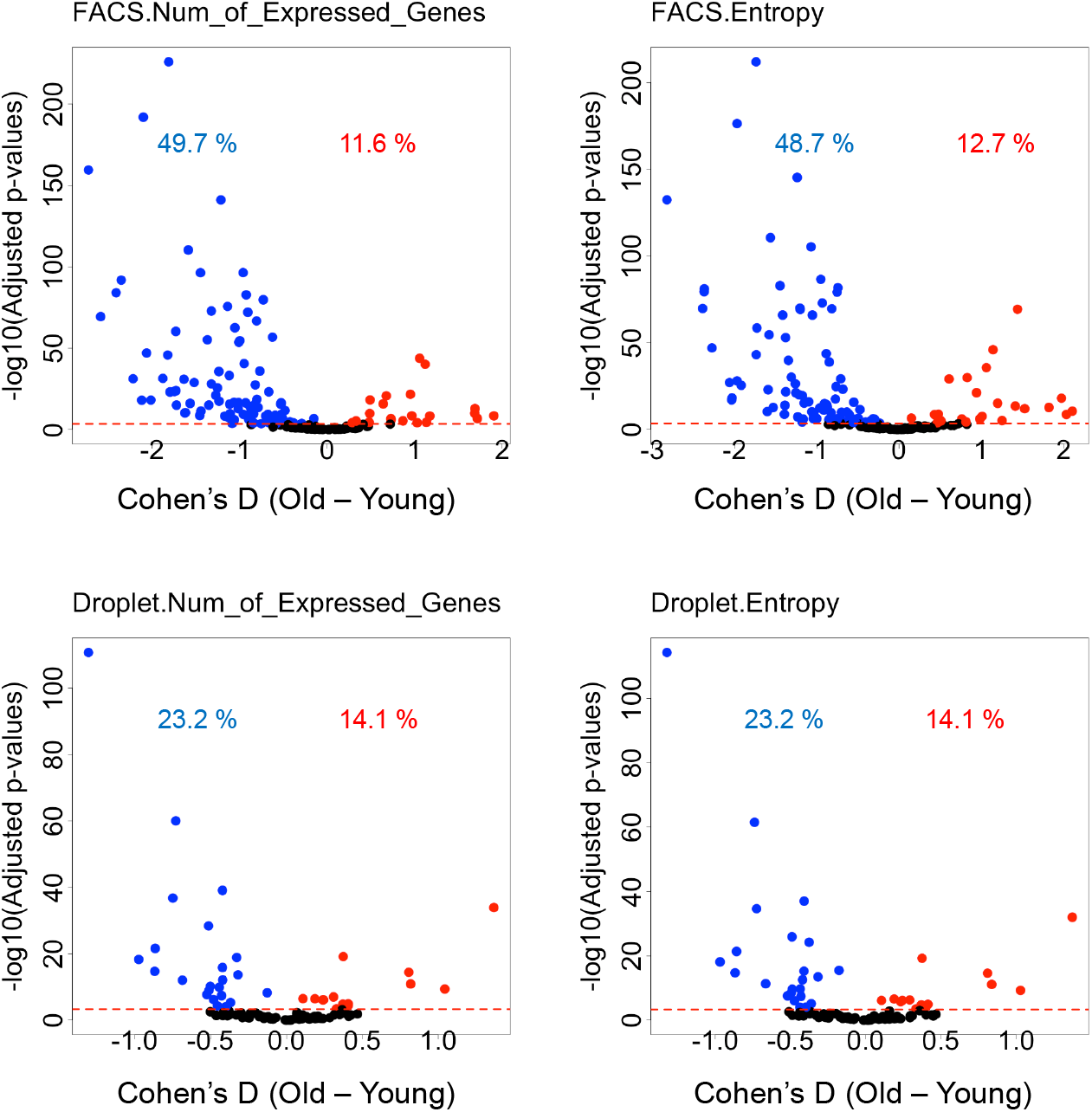
Volcano plot of age-related changes in transcriptome diversity quantified with log2(the number of expressed genes) or Shannon entropy. Each point represents one cell subset. The x-axis is the effect size of the difference between the old group and the young group as quantified by Cohen’s D. The y-axis represents −log10 (P value). Cell subsets for which statistically significant differences were detected are plotted in red (Old *>* Young) or blue (Old *<* Young) with percentages.

The results suggest that not only the direction of expression of individual genes but also the direction of aging changes in transcriptome diversity depends on the cell subset. The P-values and Cohens’D value for the analysis in each subset in the two data sets are available in Supplementary Tables S4 - S7.

## Discussion

In this study, we analyzed the Tabala Murirs Senis dataset and reported the landscape of the opposite aging effect on transcriptome profiles among cell subsets. The opposite aging effect was widely observed in the aging changes both in gene expression level and in the transcriptome diversity quantified by the number of expressed genes and the Shannon entropy.

The opposite aging effect on transcriptome can reflect a deviation from the optimal gene expression state in each cell subset. The optimal gene expression state for each cell subset is maintained in youth, but shifts to an abnormal state with age may be responsible for the dysfunctions associated with aging. The changes in gene expression associated with aging can be described as a drift in an arbitrary direction rather than an active drift in a specific direction. Gene expression is optimally regulated in each cell subset by gene regulatory networks. Each cell subset has a different steady state of gene expression due to different epigenomic states and external environments. It has been suggested that the disturbance of gene expression not only due to genetic effects by the opposite eQTL effect but also due to aging is dependent on biological conditions.

The opposite aging effect was widely observed in the genome-wide genes. In particular, the opposite aging effect enriched the genes associated with basic cellular functions such as ribosomal function and translation. It has been known that ribosomal function decreases with aging [17, 18, 21, 24]. Destabilization of the regulation of ribosome-related gene expression, represented by the oppoisite aging effect, may lead to a decrease in cellular ribosomal function.

In addition, about 30% of the known aging-related genes were observed in either of the two large data sets. If different subsets differ in the direction of aging changes in gene expression, then over-expression of the gene leads to repress aging-related changes in one cell subset but rather accelerete changes in another subset. This study highlights the importance of considering the cell subset when intervening with aging-related genes.

## Conclusion

This study reports the overall picture of the opposite aging effect, in which aging increases gene expression in one cell subset and decreases it in another cell subset. The opposite aging effect among cell subsets is widely observed in the age-related change of each gene expression level and transcriptome diversity.

## Declarations

### Ethics approval

Not applicable.

### Data and Code availability

This study used public data only. The code used in this study is available at (https://github.com/DaigoOkada/oppagingeffect).

